# Targeting primary and metastatic uveal melanoma with a G protein inhibitor

**DOI:** 10.1101/2021.01.16.426957

**Authors:** Michael D. Onken, Carol M. Makepeace, Kevin M. Kaltenbronn, Joelle Choi, Leonel Hernandez-Aya, Katherine N. Weilbaecher, Kisha D. Piggott, P. Kumar Rao, Carla M Yuede, Alethia J. Dixon, Patrick Osei-Owusu, John A. Cooper, Kendall J. Blumer

## Abstract

Uveal melanoma (UM) is the most common intraocular tumor in adults. Nearly half of UM patients develop metastatic disease and often succumb within months because effective therapy is lacking. A novel therapeutic approach has been suggested by the discovery that UM cell lines driven by mutant constitutively active Gq or G11 can be targeted by FR900359 (FR) or YM-254890, which are bioavailable, selective inhibitors of the Gq/11/14 subfamily of heterotrimeric G proteins. Here, we have addressed the therapeutic potential of FR for UM. We found that FR inhibited all oncogenic Gq/11 mutants reported in UM. FR arrested growth of all Gq/11-driven UM cell lines tested, but induced apoptosis only in a few. Similarly, FR inhibited growth of, but did not efficiently kill, UM tumor cells from biopsies of primary or metastatic tumors. FR evoked melanocytic redifferentiation of UM tumor cells with low (class 1), but not high (class 2), metastatic potential. FR administered systemically below its LD50 strongly inhibited growth of PDX-derived class 1 and class 2 UM tumors in mouse xenograft models, and reduced blood pressure transiently. FR did not regress xenografted UM tumors, or significantly affect heart rate, liver function, hematopoiesis, or behavior. These results indicated the existence of a therapeutic window in which FR can be explored for treating UM, and potentially other diseases caused by constitutively active Gq/11.

## Introduction

Mutant constitutively active forms of Gq or G11 (Gq/11) α-subunits of heterotrimeric G proteins cause uveal melanoma (UM) (1–3), and several other diseases and disorders (4–11). UM is particularly devastating because patients with metastatic disease often succumb within months due to lack of effective therapy (12). Clinical trials of metastatic UM have shown little benefit of cytotoxic chemotherapeutics, immune checkpoint inhibitors, or small molecule inhibitors of oncogenic signaling proteins including protein kinase C and MEK (13). New therapeutic strategies for metastatic UM clearly are warranted.

Recent evidence suggests that mutant constitutively active Gq/11 could be targeted therapeutically in UM and other diseases (14). Oncogenic Gq/11 α-subunits are constitutively active due to defects in hydrolyzing GTP to GDP, the rate-limiting step for G protein deactivation. Nevertheless, oncogenic Gq/11 can be inhibited by FR900359 (FR) or YM-254890 (YM) (15–20), a pair of closely related, bioavailable cyclic depsipeptides that allosterically inhibit GDP release by the Gq/11/14 subfamily (16, 21). Inhibition is thought to occur because oncogenic Gq/11 α-subunits can spontaneously release GTP, stochastically bind GDP, bind YM or FR (YM/FR) and Gβγ subunits (17), forming complexes resistant to activation by G protein-coupled receptors (14, 16) and diminishing downstream oncogenic signaling (17–20). In response to YM/FR, UM cell lines driven by oncogenic Gq/11 arrest the cell cycle, and can undergo apoptosis and/or re-differentiation into melanocytic-like cells (17–20). In contrast, BRAF-driven UM cell lines, which express wild type Gq/11, are unaffected (17–20).

The therapeutic potential of YM/FR in UM is an important opportunity based on this evidence. Critical issues and questions remain to be addressed, however. YM/FR may exert deleterious as well as therapeutically beneficial effects, because they do not discriminate between wild-type and oncogenic Gq/11 (14), and target Gq/11-dependent physiological systems (21–24) essential for homeostasis and viability (25–28). Indeed, YM/FR administered systemically at levels that reduce Gq/11 activity in host systems greater than 50% is likely to be lethal, as suggested by gene dosage studies of Gq/11 knockout mice (29). Indeed, how strongly oncogenic Gq/11 signaling in UM tumors must be inhibited for therapeutic effect remains unknown. Furthermore, although recent studies have shown that YM/FR can inhibit growth of UM tumors in mouse xenograft models (19, 30), left unanswered are whether YM/FR can target primary and/or metastatic UM tumor cells from patients, whether YM/FR responsiveness is affected by tumor-intrinsic factors, including the type of oncogenic Gq/11 mutation, tumor class or metastatic potential, or whether systemic administration of YM/FR at levels sufficient to target UM tumor xenografts has serious deleterious effects on viability or host physiological systems. Here, we have addressed all of these important issues.

## Results

### FR inhibits all oncogenic Gq and G11 mutants that drive uveal melanoma

At least 10 different constitutively active Gq/11 mutants have been identified in UM (2, 31, 32). Only four had been studied previously and shown to be inhibited by FR or YM (17–20). To extend this analysis, we transiently expressed each of ten different Gq/11 mutants that drive UM (Fig. 1A) with a Gq/11-driven transcriptional reporter in a common cellular context (HEK293 cells) to exclude extrinsic factors that might affect the results. We found that each mutant G protein induced reporter expression and was inhibited by FR with similar potency (IC_50_ 1.9 to 3.8 nM) and efficacy (Fig. 1A).

**Fig. 1:**
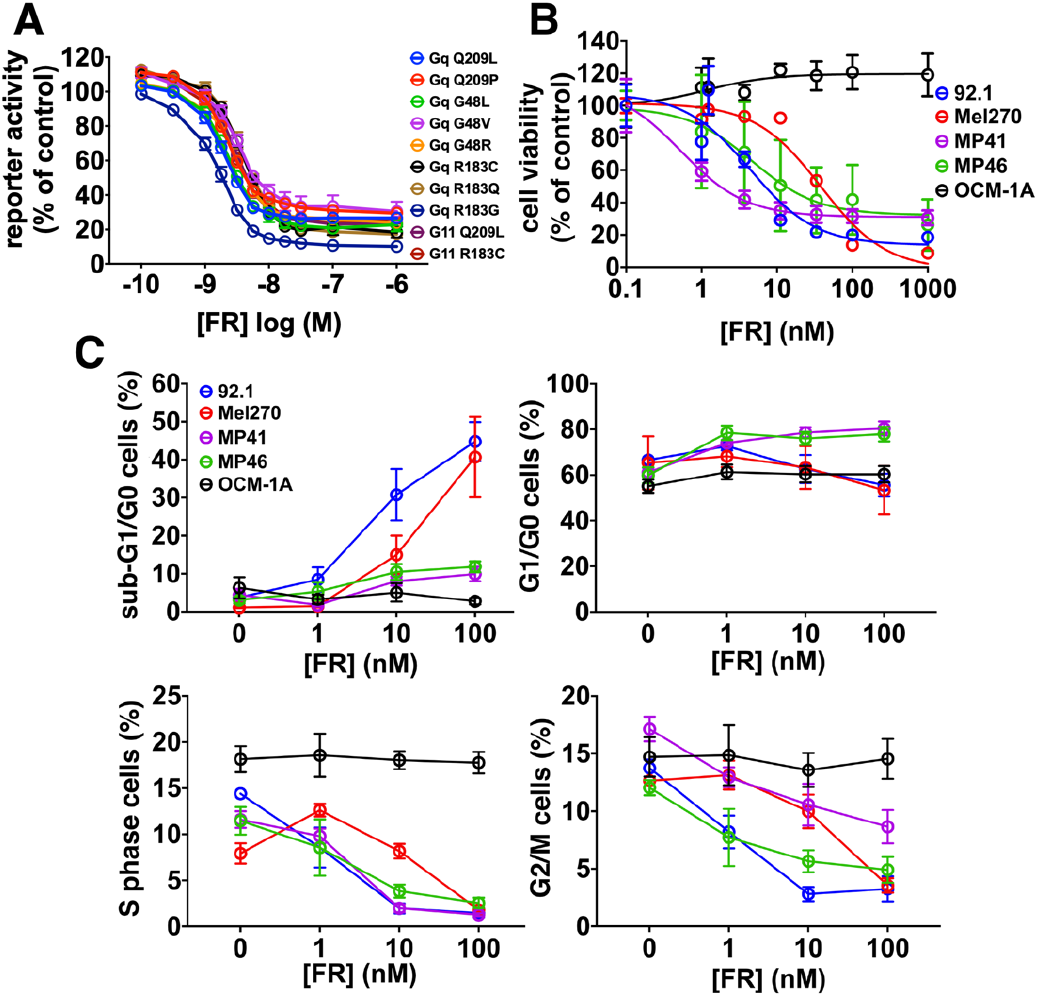
Effects of FR on constitutively active Gq/11 mutants and UM cell lines. A) Effects of FR on expression of an SRE.L reporter driven by transfection of oncogenic Gq/11 mutants in HEK293 cells. FR inhibited all 10 Gq/11 mutants reported in UM, with IC50s between 1.9 nM (Gq-Q209L) and 3.8 nM (Gq-R183Q). B) Effects of FR on UM cell lines. Growth of UM cell lines driven by constitutively active mutants of Gq or G11 was inhibited by FR with IC_50_s between 0.5 nM (MP41) and 38 nM (Mel270). A UM cell line (OCM-1A) driven by constitutively active BRAF(V600E) was unaffected by FR. C) Effect of FR on apoptosis and cell cycle progression of UM cell lines detected by flow cytometry. Induction of apoptosis was indicated by increased proportion of cells with sub-G1/G0 DNA content. Cell cycle arrest was indicated by decreased proportion of cells with G1/G0, S-phase and G2/M-phase DNA content.

In contrast, we found that four UM cell lines (Fig. 1B) driven by three different oncogenic Gq/11 mutants responded to FR in quantitatively or qualitatively different ways. FR inhibited growth of all four Gq/11-driven UM cell lines, but with potencies differing by as much as ~80-fold (IC_50_ 0.5 nM (MP41 cells), 38 nM (Mel270 cells); Fig. 1B). Whereas FR arrested the cell cycle of all Gq/11-driven UM cell lines, as indicated by reduction of S- and G2/M-phase cells (Fig.1C and Fig. S1), it induced significant apoptosis only in two of them (92.1 and Mel270 cells), as indicated by accumulation of cells with sub-G0/G1 DNA content (Fig. 1C and Fig. S1). FR did not affect proliferation or survival of a BRAF(V600E)-driven UM cell line (Fig. 1B), as shown before (17–19), demonstrating the selectivity of FR for oncogenic Gq/11 in UM.

### FR re-differentiates class 1 but not class 2 UM cells

Class 1 and class 2 primary UM tumors have low (~5%) and high (~90%) probability of metastasis, respectively (33, 34). Class 1 UM tumors express the metastasis suppressor, BAP1, encoding an enzyme that regulates gene expression epigenetically by deubiquitinating histone H2A. Class 2 UM tumors are BAP1-deficient (35) and possess additional genetic defects rendering them prone to metastasis (32). Almost all UM cell lines are derived from class 1 tumors (36), raising the crucial question of whether both classes of primary UM tumor cells can be targeted by FR.

We addressed this question initially by analyzing MP41 (class 1; BAP1^+^) and MP46 (class 2; BAP1-deficient) cell lines, which were established originally from patient-derived xenografts (PDX) of primary UM tumors (37). We found that FR arrested growth and cell cycle progression of both of these cell lines without triggering significant apoptosis (Fig. 1, B and C).

Next, we used transcriptional analysis to determine whether FR could re-differentiate MP41 (class 1) and MP46 (class 2) cells into melanocytic-like cells, as we had shown previously with two other class 1 UM cell lines (92.1 and Mel202) (17). MP41 and MP46 cells both responded transcriptionally to FR, as indicated by multi-dimensional scaling analysis of RNAseq data (Fig. 2A). However, a cluster of genes was downregulated preferentially in class 1 MP41 cells (circled in Fig. 2B), as revealed by analyzing transcriptional responses on a gene-by-gene basis. Many genes in this cluster are targets of epigenetic regulation by polycomb repressive complex 2 (PRC2), based on gene set analysis (H3K27trimethylation, EZH2; Fig. 2C). This PRC2-targeted gene cluster was coordinately repressed by FR in class 1 MP41 cells as compared to class 2 MP46 cells (Fig. 2D). Consistent with these distinctions, FR induced pigmentation, a hallmark of melanocytic differentiation, of MP41 but not MP46 cells (Fig. 2D). Indeed, our prior studies had shown that PRC2 activity is required for FR to reinstate melanocytic differentiation of two other class 1 UM cell lines (92.1 and Mel202) (17). Therefore, the ability of FR to promote melanocytic redifferentiation of primary UM tumors may depend on tumor class.

**Fig. 2:**
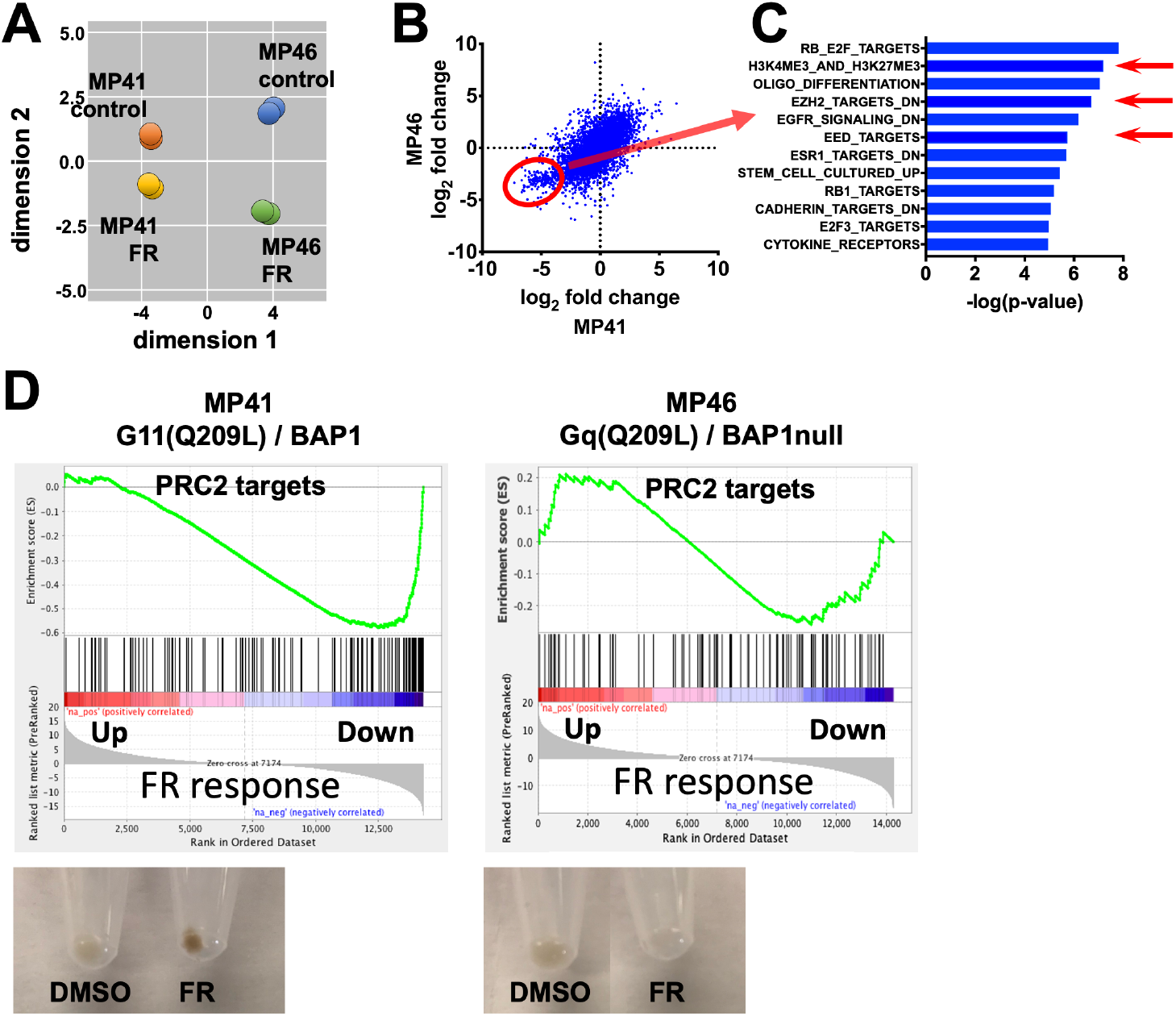
Transcriptional responses elicited by FR in UM cell lines that model class 1 and class 2 tumors. A) Multidimensional scaling plot of RNAseq data showing similar directional responses of MP41 (class 1) and MP46 (class 2) cells to FR. B) Scatter plot comparing the effects of FR on gene-by-gene basis in MP41 and MP46 cells. Each blue dot represents a single gene. Genes circled in red were downregulated by FR significantly more strongly in MP41 cells relative to MP46 cells. C) Gene set enrichment analysis of genes preferentially downregulated by FR in MP41 cells. Red arrows indicate PRC2-targeted gene sets. D) Gene set enrichment analysis indicating the effects of FR on expression of PRC2-targeted genes in MP41 and MP46 cells. Images of pelleted cells (below) indicating the effects of FR on pigmentation as a marker of melanocytic differentiation of MP41 and MP46 cells.

### FR targets UM cells from primary and metastatic tumors

Because UM tumor cell lines do not fully recapitulate important properties of UM tumors, we determined whether FR can target UM cells obtained from patient biopsies of primary and metastatic tumors. We began by preparing self-renewing cultures of UM tumor cells isolated from fine-needle biopsies of class 1 primary tumors from six patients, and core biopsies of liver metastases from two patients. We found that all eight cultured-cell samples responded to FR, as indicated by lower cell numbers over time relative to vehicle controls (Fig. 3A), and we noted, as shown before (36, 37), that class 2 primary UM tumor samples proliferated too poorly in vitro to establish self-renewing cultures.

**Fig. 3:**
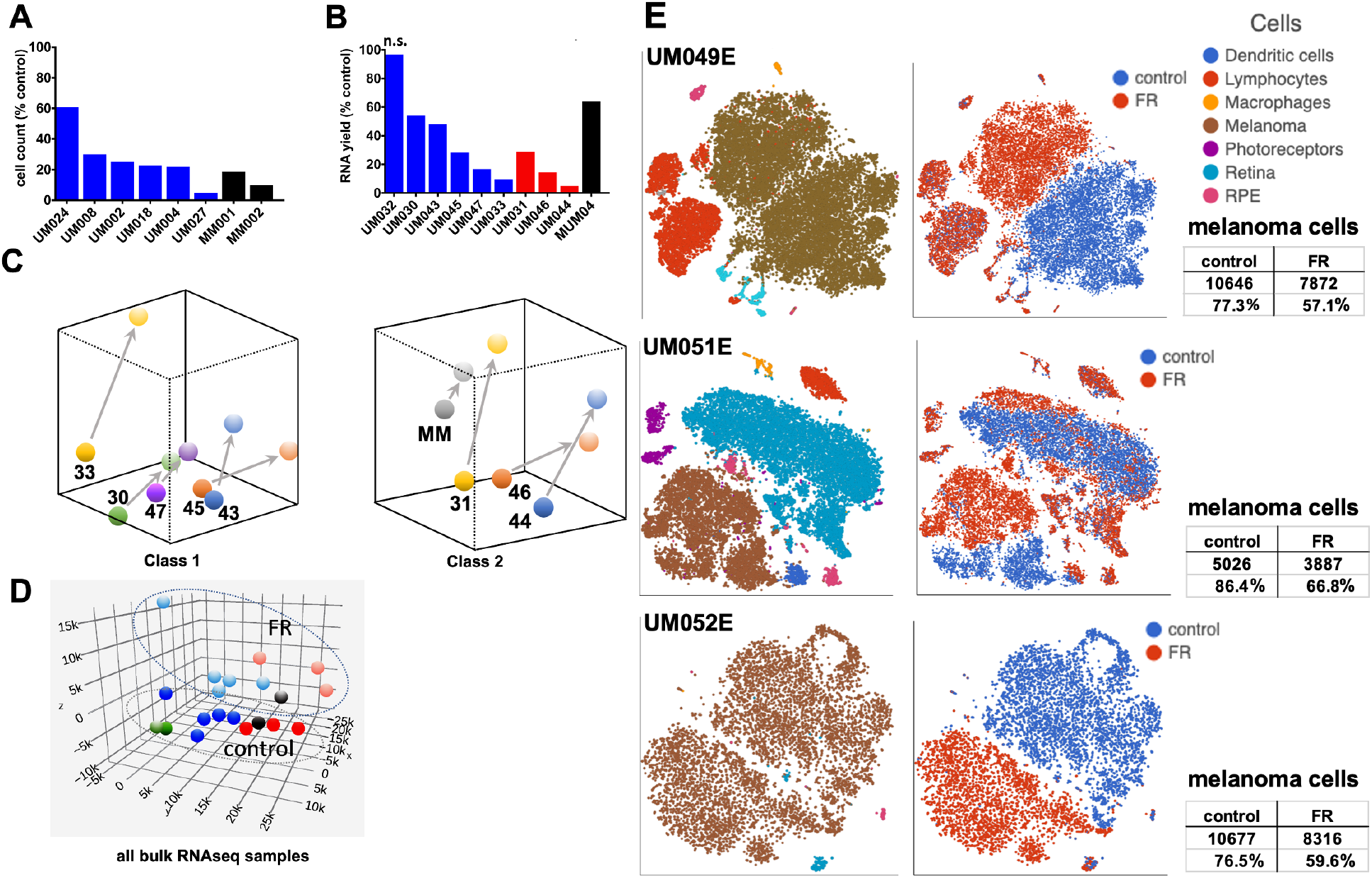
Response of human UM tumor biopsy samples to FR ex vivo. A) Preliminary set of human tumor fine-needle biopsy samples were established in culture and then split equally. Cells were counted after 7 d of culture in absence or presence of FR (100 nM). B) Human biopsy samples from class 1 and class 2 primary UM tumors and a liver metastasis were dissociated, split equally and treated immediately for 7 d with vehicle or FR (100 nM). The effect of FR relative to the vehicle control on the yield of total RNA as a marker of cell number is shown. One sample (UM032) expressed wildtype Gq and G11 and did not respond to FR. C) PCA plots of RNAseq data from class 1 (left) and class 2 (right) human tumor biopsy samples. Arrows indicate the directional effect of FR on gene expression for each sample. D) PCA plot of all samples combined. Class 1 samples are blue; class 2 samples are red; metastatic sample is gray; and unresponsive sample (UM032) is green. E) 2-dimensional t-SNE plots of single-cell scRNAseq data for each of three primary UM tumors obtained after enucleation, treated 7 d with vehicle or FR (100 nM). Left panels are color coded by cell type; right panels are color coded by treatment. Non-melanoma cell clusters (left panels) show overlap of red and blue cells (right panels), suggesting minimal transcriptional response to FR. Melanoma cells are segregated in FR-(red) and vehicle-treated (blue) samples (right panels), indicating marked differences in gene expression.

Accordingly, we switched to short-term cultures of biopsy samples obtained from ten additional patients (Table S1) that included class 1 and class 2 primary tumors and a liver metastasis. Cultured cells from nine of these patient samples responded to FR relative to vehicle controls, as indicated by decreased levels of total RNA as a marker of cell number (Fig. 3B). The single non-responding sample (UM032; Fig. 3B) expressed wild type Gq and G11 (Table S1), and we excluded it from subsequent studies of FR-induced transcriptional responses. For the nine FR-responsive UM tumor samples, which included class 1 and class 2 primary tumors and a liver metastasis, we found similar directional effects on gene expression, as revealed by unsupervised principal component analysis (PCA) (Fig. 3C). The principal component representing transcriptional differences between class 1 and class 2 tumor cells (Fig. 3D; x-axis) (33) distinguished both vehicle- and FR-treated cells (Fig. 3D), demonstrating that FR did not cause class 2 UM tumor samples to lose their class identity and become more class 1-like. Together, these results demonstrated that, regardless of tumor class, primary and metastatic UM tumor samples driven by mutant Gq/11 respond ex vivo to FR.

Because patient-derived UM tumor biopsies include cancer and stromal cells, we determined which cell types were targeted by FR, as indicated by their relative abundance and transcriptional responses revealed by single-cell RNAseq (scRNAseq). Primary tumor samples from three UM patients undergoing enucleation were used to obtain sufficient quantity of cells for these experiments. Transcriptional signatures readily distinguished melanoma cells from several stromal cell types (Fig. 3E). Dissociated UM tumor and stromal cells were treated 7 days with FR at a concentration (100nM) that strongly inhibits all oncogenic Gq/11 mutants (Fig. 1A). In all three biopsy samples, FR modestly reduced the relative abundance of melanoma cells (Fig. 3E), indicating that extensive tumor cell death had not occurred.

Further analysis of scRNAseq data indicated that melanoma cells, but not stromal cells, displayed striking transcriptional responses to FR (Fig. 3E). Melanoma cells in all three biopsy samples showed directional changes in gene expression (Fig. 3E) analogous to those observed previously by bulk RNAseq of small biopsy samples (Fig. 3D). FR-evoked gene expression responses detected by scRNAseq of enucleated tumors and bulk RNAseq of small biopsies were similar, as shown by hallmark enrichment analysis (Fig. 4A). In both cases, targets of E2F transcription factors (arrows in Fig. 4, A and B) were among the gene sets prominently downregulated by FR in both class 1 and class 2 tumor biopsy samples. Moreover, FR treatment of class 1 and class 2 biopsy samples downregulated expression of the E2F transcription factors E2F1, E2F2 and E2F8 (Fig. 4C), and upregulated expression of the cell cycle inhibitors RB1 and p130 (38). In sum, inhibition of oncogenic Gq/11 by FR upregulated Rb expression and downregulated E2F expression, thereby contributing to inhibition of cell cycle progression (Fig. 4D).

**Fig. 4:**
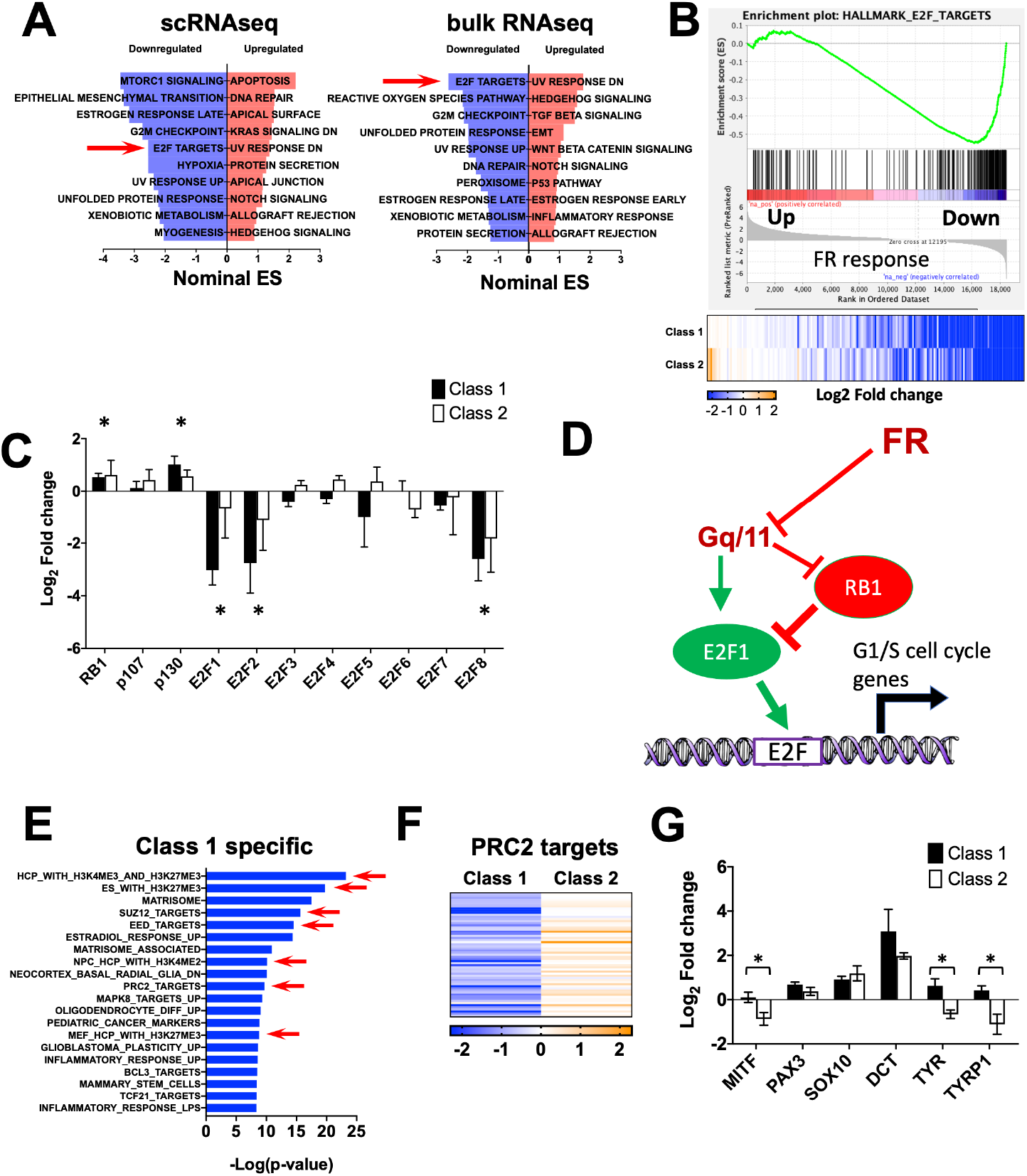
Pathway analysis of FR transcriptional responses. A) Hallmark gene-set enrichment analysis of differential gene expression in response to FR. Left panel shows results from melanoma cells annotated from scRNAseq data. Right panel shows analysis of bulk RNAseq data from 9 biopsy samples described in Figure 3. Red arrow highlights E2F targets significantly downregulated in both scRNAseq and bulk RNAseq datasets. B) Enrichment plot of E2F target hallmark gene set in bulk RNAseq data with heatmap below showing no significant differences in response in class 1 versus class 2 biopsy samples. C) Effects of FR on expression of Rb/E2F family members in class 1 and class 2 biopsy samples detected by RNAseq. Asterisks indicate p < 0.01 relative to vehicle controls. Differences between class 1 and class 2 samples were insignificant. D) Proposed model for cell cycle regulation by constitutively active Gq/11 and FR. Signaling by constitutively active Gq/11 leads to repression of Rb expression and induction of E2F expression such that FR attenuates these effects. E) Differential responses of class 1 and class 2 tumor biopsy samples to FR. Gene set enrichment analysis identifies genes downregulated in class 1 but not class 2 tumor biopsy samples. Red arrows indicate PRC2-specific gene sets. F) Heat map showing effects of FR on PRC2-targeted genes in class 1 versus class 2 tumor samples. G) Responses of melanocytic differentiation genes to FR in class 1 and class 2 tumor biopsy samples. Asterisks indicate statistically different (p < 0.01) effects of FR on the indicated melanocytic differentiation genes in class 1 and class 2 biopsy samples.

Next, we investigated whether tumor class, which predicts metastatic potential and patient prognosis, determines whether FR evokes melanocytic redifferentiation of UM tumor cells in patient biopsies, as we had observed above with UM cell lines (Fig. 2). When we examined transcriptional responses of class 1 UM tumor cells to FR, we found strong enrichment of gene sets targeted by PRC2 (Fig. 4E). Nearly all members of the FR-responsive PRC2-targeted gene cluster identified in UM cell lines (Fig. 2) were also downregulated in class 1 UM tumor biopsies (Fig. 4F). In contrast, many of these genes were upregulated, rather than downregulated by FR in class 2 UM tumor biopsies (Fig. 4F). Furthermore, melanocyte differentiation genes, including MITF, TYR, and TYRP1, were upregulated significantly by FR in class 1 relative to class 2 tumor biopsy samples (Fig. 4G). Therefore, FR treatment potentially could inhibit progression of class 1 primary UM tumors by promoting melanocytic redifferentiation.

### FR up- and down-regulates YAP1-driven genes

Prior studies of UM have shown that oncogenic Gq/11 signaling leads to activation of the YAP1 transcription factor (19, 20). We therefore determined whether YAP-targeted genes are coordinately downregulated by FR in biopsy samples of class 1 and class 2 primary UM tumors. In contrast to this expectation, we found that FR upregulated and downregulated YAP-targeted genes in nearly equal proportion (Fig. S2). The mechanisms responsible for the complex effects of FR on YAP-targeted genes are unknown, but potentially involve differential regulation by other transcription factors in various combinations with YAP1 on the promoters of these genes.

### A therapeutic window for targeting UM tumors with FR

Because inhibition of Gq/11 activity by more than 50% is likely to be lethal, as indicated by gene dosage studies in knockout mice (29), we sought to define protocols and conditions under which chronic, systemic administration of FR is physiologically tolerated, so that we could assess the therapeutic potential of FR in mouse xenograft models of UM. Based on prior studies examining the acute effects of FR or the closely related inhibitor (YM-254890 (YM)) (22, 24, 39), we administered FR by subcutaneous injection at 0.1-3.0 mg/kg on alternate days for 30 days to define an LD50 of ~0.6 mg/kg in NOD-scid-gamma (NSG) mice that were used as xenograft recipients (Fig. 5A). We then determined the effect of FR administered chronically below its LD50 (0.1 or 0.3 mg/kg by subcutaneous injection on alternate days) on several physiological parameters of NSG mice. Blood pressure, a sensitive indicator of Gq/11 activity (40), decreased significantly in conscious mice upon each injection of FR relative to vehicle controls (Fig. 5B), but rebounded to baseline within 24 h (Fig. 5C). Heart rate was affected insignificantly by FR (Fig. 5B). No evidence of anemia, monocyte or platelet deficiency, or liver disfunction was observed after treating NSG mice 30 d with FR at either dose relative to vehicle controls (Fig. 5D). Lastly, even when administered at the higher dose, FR had insignificant effect on the activity, exploratory behavior, or sensorimotor function of NSG mice (Table 1 and Fig. S3), even though Gq/11 are important regulators of nervous system development and activity (40). Chronic systemic administration of FR below its LD50 therefore had surprisingly modest physiological effects that potentially could limit dosing regimens in clinical trials.

**Fig. 5:**
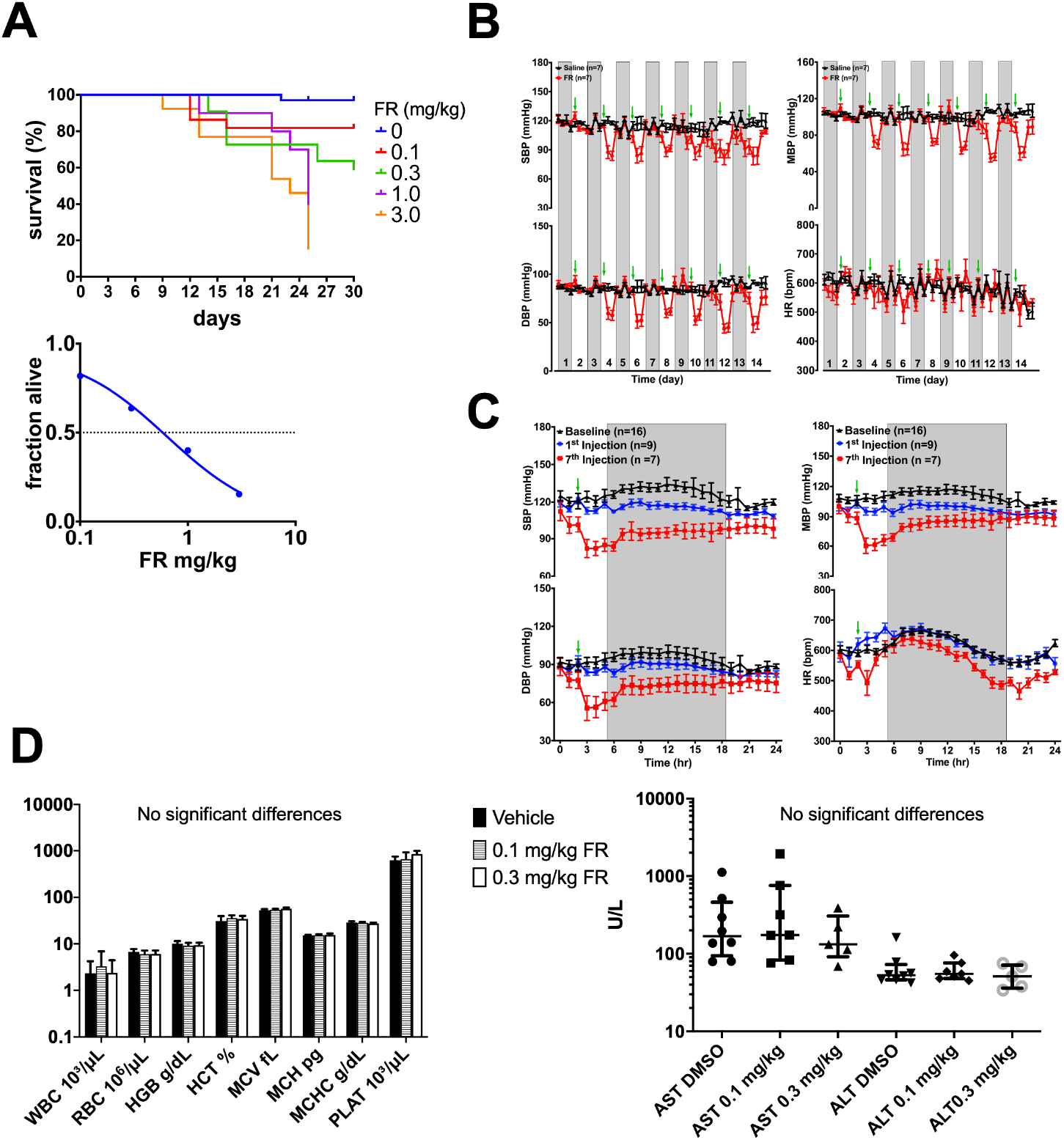
Effects of systemically administered FR on mouse viability and physiology. A) Survival analysis of NSG mice treated up to 30 days with FR administered at the indicted doses by s.c. injection on alternate datys. Kaplan-Meier (upper panel) and survival (lower panel) curves indicate an LD50 of ~ 0.6mg/kg. B) Effect of FR on blood pressure as an indicator of Gq/11 inhibition in host tissues. Adult NSG male mice received subcutaneous administration of vehicle (2 μl DMSO in 200 μl of 5% dextrose, saline; n=7) or FR (0.3 mg/kg, n=9) once every other day for 14 d. Blood pressure and heart rate were monitored telemetrically 1 h before and 3 h after injection. The injection times are indicated by green arrows. Days without radiotelemetry recording or injection are indicated by gray columns. C) Twenty-four-hour continuous radiotelemetry recordings of blood pressure and heart rate in mice before the start of the injection regimen (Baseline, black line, n=16), and after the 1^st^ (blue line, n=9) and 7^th^ injections (red line, n=7), respectively. The shaded regions represent 12-h dark periods, while the green arrows indicate the time of injection. All values are mean ± SEM. Blood pressure (BP) and heart rate (HR) values are plotted at 1 h intervals. SBP, systolic blood pressure; DBP, diastolic blood pressure; MBP, mean blood pressure; HR, heart rate. D) Effect of FR on hematopoiesis and liver function. Blood samples were collected from mice after 30 d of treatment with FR (0.3 mg/kg s.c. on alternate days) and assayed for blood cell counts (left panel) and liver enzyme activities (right panel). No significant changes were seen with FR treatment.

**Table 1:** Effect of FR on mouse behavior

| Variable | CTL mean | TX mean | Diff ± SEM | P Value |
| --- | --- | --- | --- | --- |
| Walking initiation (s) | 2.54 | 1.98 | 0.96 ± 1.57 | p = 0.3537 |
| Ledge (/60s) | 48.92 | 43.35 | 29.12 ± 19.79 | p = 0.4315 |
| Platform (/60s) | 22.97 | 25.74 | 4.4 ± 18.55 | p = 0.7294 |
| Pole (/120s) | 25.92 | 22.16 | 0.84 ± 25.07 | p = 0.5844 |
| Turn (/60s) | 23.73 | 15.97 | -1.79 ± 25.52 | p = 0.5844 |
| 60 degree screen (/60s) | 11.43 | 10.68 | 7.88 ± 3.54 | p = 0.634 |
| 90 degree screen (/60s) | 16.98 | 16.49 | 7.04 ± 9.93 | p = 0.9049 |
| Inverted screen (/60s) | 49.92 | 41.10 | 32.17 ± 17.74 | p = 0.1437 |
| Fine Movements (/1h) | 1758 | 1549.00 | 209.1 ± 115.5 | p = 0.0861 |
| Total Ambulations (/1h) | 756.5 | 674.3 | -82.20 ± 64.16 | p = 0.2155 |
| Time Rearing (/hr) | 688.5 | 583.3 | 105.1 ± 81.22 | p = 0.2111 |
| Center Distance (mm/h) | 4288 | 3941 | 347.4 ± 506.5 | p = 0.5010 |
| Center Time (/1h) | 478.3 | 557.6 | -79.30 ± 112.8 | p = 0.4905 |
| Center Entries (/1h) | 263.9 | 236.1 | 27.81 ± 27.70 | p = 0.3281 |
| Total Rest Time (s/1hr) | 1504 | 1635 | -130.6 ± 118.7 | p = 0.2847 |
| Total Ambulations over time | 4539 | 4046 | 493.2 ± 385.0 | p = 0.2733 |
| Rearing frequency over time | 116.8 | 123.5 | 6.671 ± 14.51 | p = 0.8508 |

Next, we determined whether FR delivered systemically at physiologically tolerated doses could inhibit growth of xenografted UM tumors driven specifically by oncogenic Gq/11. These experiments used MP41 and MP46 cells for several reasons. They were isolated originally from patient-derived xenografts, provide models of class 1 and class 2 tumors, and, of the UM cell lines tested, responded to FR (cell cycle arrest, with little or no apoptosis) most like UM tumor cells from patient biopsies. We also used a UM cell line (OCM-1A) driven by BRAF(V600E) to determine whether any effects of FR were specific for Gq/11-driven tumors. Accordingly, subcutaneous UM tumor xenografts of MP41, MP46 and OCM-1A cells were established, and vehicle or FR was administered systemically at low (0.1 mg/kg) or high (0.3 mg/kg) dose on alternate days by subcutaneous injection on the contralateral side. Tumor size was measured over time and tumor weight was determined at the end of the experiment. Relative to vehicle controls, overall survival of these tumor-bearing cohorts of mice treated ~30 d with FR at low or high dose was, respectively 85% and 71%. Under these conditions, FR inhibited growth of MP41 tumors only at the high dose (Fig. 6A), but inhibited growth of MP46 tumors at either dose (Fig. 6A). FR had no effect on growth of BRAF(V600E)-driven OCM-1A tumors (Fig. 6A), demonstrating that FR did not affect processes such as tumor vascularization or perfusion that generically support tumor growth. Thus, FR specifically targeted both class 1 and 2 UM tumors, but only those driven specifically by oncogenic Gq/11.

**Fig. 6:**
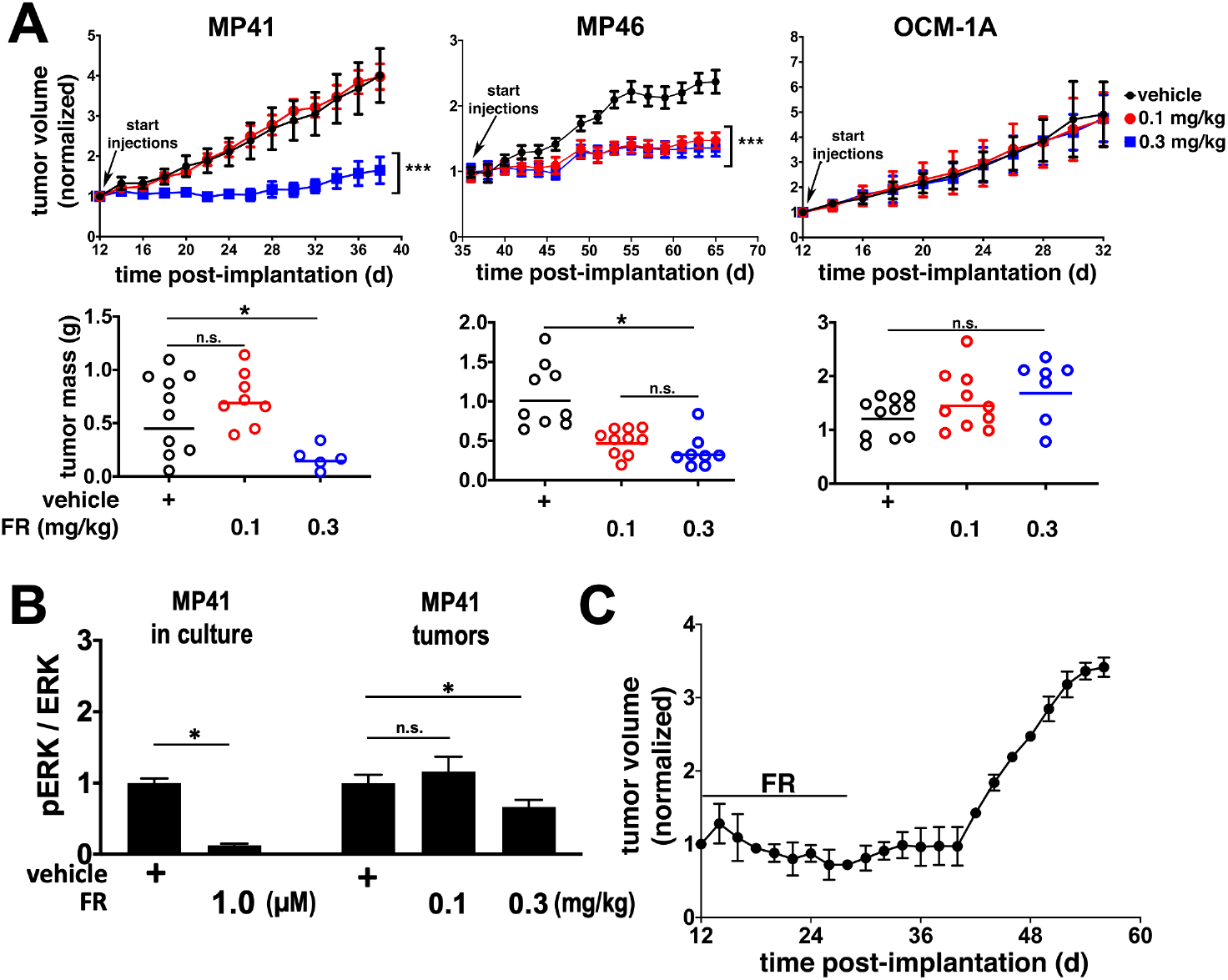
FR inhibits growth of xenografted class 1 and class 2 UM tumors. A) Tumor volume (upper panels) as a function of time and dose of FR administered at the indicated dose by s.c. injection on alternate days. Effect of FR on tumor weight (lower panels) measured at the end of the experiment. B) Effect of FR on Erk phosphorylation as a marker of oncogenic G11 activity in MP41 cells in vitro versus tumors from animals treated with vehicle or FR at the indicated doses. Shown are the effect of FR on pERK:total Erk ratios relative to vehicle controls. C) Durability of FR response by MP41 mouse xenograft tumors treated systemically with FR for 25 d, after which time FR treatment was stopped. Tumors resumed growth 16 d after FR treatment was withdrawn. Asterisks indicate p < 0.05 for panels A and B.

To determine whether FR delivered systemically at therapeutically effective doses inhibited oncogenic G protein signaling in vivo, we compared the effects of FR on Erk activity, one of several downstream pathways activated by oncogenic Gq/11 in UM (41), in vivo vs. in vitro (Figs. 6B and S4). FR administered at a therapeutically effective dose (0.3 mg/kg on alternate days) reduced Erk phosphorylation in G11-driven MP41 tumors in vivo by ~30%. In contrast, Erk phosphorylation in MP41 cells in vitro was inhibited strongly (85%) at a high concentration of FR. Thus, modulation rather than complete suppression of oncogenic G protein signaling in UM tumor xenografts was sufficient for FR to have therapeutic effect.

Lastly, we investigated whether FR caused durable arrest or regression of class 1 UM tumor xenografts. In this experiment, mice were implanted with MP41 cells and allowed to form tumors. Tumor growth was arrested by treating animals with FR at an effective dose (0.3mg/kg s.c. on alternate days) for 25 d (Fig 6C). FR treatment then was stopped. Tumors resumed growth 16 d later (Fig. 6C). Therefore, FR appeared to cause reversible rather than durable arrest or regression UM tumors in this model. Taken together, these results indicated the existence of a therapeutic window in which FR effectively targeted class 1 or class 2 UM tumor xenografts, while preserving sufficient function of Gq/11 in host tissues required to support essential physiological processes and viability.

## Discussion

### Targeting UM tumors with FR

Here we have provided several new lines of evidence that support the therapeutic potential of Gq/11 inhibitors such as FR for UM. Our findings indicate that the vast majority of primary and metastatic UM tumors potentially can be targeted by FR. We found that all constitutively active mutant forms of Gq/11 found in UM, which collectively drive tumorigenesis in >90% of patients, are sensitive to FR. Moreover, FR can target Gq/11-driven UM tumor cells from patient biopsies of primary ocular tumors with low or high metastatic potential, and liver metastases. And lastly, although FR does not discriminate between wild type and oncogenic Gq/11, and host Gq/11 activity is physiologically essential, we were able to identify a therapeutic window in which FR strongly inhibited tumor growth without large negative effects on viability or physiology. UM tumors therefore appear to be particularly vulnerable to FR.

The vulnerability of UM tumors to FR may occur because the signaling networks downstream of oncogenic Gq/11 must be maintained above a high threshold. In support of this hypothesis, we found that a therapeutically effective dose of FR strongly inhibits UM tumor growth but only modestly reduces Gq/11-driven Erk phosphorylation in UM tumors. Because nearly complete inhibition of Erk phosphorylation by FR was not necessary to inhibit UM tumor growth, Erk likely cooperates with other signaling pathways downstream of oncogenic Gq/11 (42) to sustain UM tumors. Indeed, Gq/11 and MEK inhibitors recently have been shown to synergize and regress UM tumor xenografts (30), suggesting that synergism might improve efficacy or safety of Gq/11 inhibitors such as FR as therapeutic agents for UM.

### Therapeutic implications

Our results suggest several ways that Gq/11 inhibitors such as FR could be explored therapeutically to treat UM patients, either alone or in conjunction with current standards of care. Primary UM tumors are usually treated by external beam or plaque irradiation of the eye (12) to arrest growth, induce tumor cell death, and preserve vision, although these interventions on their own do not improve the prognosis for metastases (13). Once metastatic UM is detected in the liver, therapy involves mechanical isolation of the hepatic vasculature by embolization and percutaneous infusion to deliver high-dose chemotherapy (43) or radiotherapy (44) above what would be tolerated systemically (45). Thus, targeting delivery of FR to primary UM tumors by intravitreal injection, or hepatic metastases by chemoembolization, could be considered. Such focal delivery could reduce the impact of FR on blood pressure or other dose-limiting physiological processes, or allow FR to be delivered at a higher dose in an effort to promote durable arrest or tumor regression. If FR proves to have limited efficacy in monotherapy, it could be combined with standard-of-care interventions to treat primary and metastatic UM. Furthermore, intravitreal injection of FR might interfere with progression of very early-stage lesions to class 2 tumors that have high metastatic potential, by promoting melanocytic differentiation, as we have shown for class 1 tumors. Similarly, because intraocular nevi are relatively common, express constitutively active Gq/11, and can progress to early stage melanomas (46), intravitreal injection of FR might interfere with this process and reduce the incidence of UM.

UM is only one of several diseases that could benefit from targeted treatment with FR. Constitutively active mutant forms of Gq/11 occur in ~5% of all cancers including melanocytic neoplasms of the central nervous system (4), mucosal melanoma (5), choroidal hemangiomas (8, 11), and hepatic small vessel neoplasms (6), as well as in Sturge-Weber syndrome (7), certain forms of hypercalcemia and hypocalcemia (9), and autosomal dominant hypoparathyroidism (10). Treatment regimens for all of these conditions potentially could be improved by novel therapeutic approaches such as FR.

## Experimental Procedures

### Reagents

FR900359 (FR) was purified from *Ardisia crenata* according to published methods (16). SRE.L assays use a firefly luciferase reporter driven by the Gq/11-dependent SRE promoter and were performed as described (47). Firefly luciferase activity was normalized to Renilla luciferase expressed from a constitutive promoter.

### Cell culture assays

Cells were cultured at 37°C in 5% CO2. Human 92.1 (RRID:CVCL_8607), Mel270 (RRID:CVCL_C302), and OCM-1A (RRID:CVCL_6934) UM cells were derived by and the generous gifts of Drs. Martine Jager (Laboratory of Ophthalmology, Leiden University), Bruce Ksander (Schepens Eye Institute, Massachusetts Eye and Ear Infirmary) and June Kan-Mitchell (Biological Sciences, University of Texas at El Paso), respectively. The human PDX cell lines MP41 (ATCC Cat# CRL-3297, RRID:CVCL_4D12) and MP46 (ATCC Cat# CRL-3298, RRID:CVCL_4D13) cell lines were purchased from ATCC (Manassas, VA). Cell lines were grown in RPMI 1640 medium (Life Technologies, Carlsbad, CA) supplemented with FBS and antibiotics. Cell viability was measured using a water-soluble tetrazolium salt, WTS-8 (Bimake, Houston, TX), following the manufacturer’s protocol. Flow cytometry for analysis of cell proliferation and apoptosis was performed at the Siteman Cancer Center Flow Cytometry Core on a FACScan analyzer (BD Biosciences, San Diego CA, USA) using a standard propidium iodide staining protocol as described previously (48).

### Patients and sample collection

Human tissue samples were obtained with patient written informed consent and with approval of the institutional review board of Washington University in St. Louis. Fine-needle aspiration biopsies of primary UM tumors and ultrasound-guided core biopsies of UM liver metastases were performed as part of standard of care, which also collected biopsies for cytological evaluation of tumor cells and molecular classification (Castle Biosciences). For enucleation samples, after excision, eyes were transilluminated to localize the tumor mass and then opened opposite the tumor. Vitreous was carefully removed and tumor samples were collected through the retina from the tumor apex. All samples were collected directly into growth medium in the OR before being transported to the laboratory. Primary uveal melanoma cells were equally divided onto two wells of a fibronectin-covered 6-well tissue culture plate and grown in 5% CO2 in MDMF medium which consists of HAM’s F12 (Lonza, Walkersville MD, USA) supplemented with 1 mg/mL BSA (Sigma-Aldrich, St Louis MO, USA), 2 mM L-glutamine (Lonza), 1X SITE (Sigma-Aldrich), 1x B27 (Gibco, Carlsbad CA, USA), 20 ng/mL bFGF (PeproTech Inc, Rocky Hill NJ, USA), and 50 μg/mL Gentamicin (Sigma-Aldrich) (49). Cells were allowed to attach to the substrate overnight before fresh medium was added to each well containing either 100 nM FR or an equal volume of vehicle (DMSO).

### Bulk RNAseq

MP41 and MP46 cells were treated with 100 nM FR or vehicle (DMSO) in RPMI growth medium and collected after 3 d of treatment. FNAB samples were treated with 100 nM FR or vehicle in MDMF medium and collected after 7 d of treatment. RNA was isolated using the RNeasy Mini Kit (Qiagen) following the manufacturer’s protocol and including the optional DNase I treatment step. RNA quality was assessed on a Bioanalyzer 2100 (Agilent Technologies, Santa Clara, CA, USA). mRNA was extracted from total RNA using a Dynal mRNA Direct kit, fragmented and reverse transcribed to double-stranded cDNA with random primers before addition of adapters for library preparation. Library preparation and HiSeq2500 sequencing were performed by the Washington University Genome Technology Access Center (gtac.wustl.edu). FastQ files were aligned to the transcriptome and the whole-genome with STAR. Biologic replicates were simultaneously analyzed by edgeR and Sailfish analyses of gene-level/exon-level features. Unexpressed genes and exons were removed from the analyses. Unsupervised principal component analysis was generated in Bioconductor using edgeR. Direct comparison of FR response in MP41 versus MP46 cells was used to identify MP41-specific genes and this list was used as signature gene sets for Gene Set Enrichment Analysis (GSEA) (50).

### Single-cell RNAseq

Single-cell RNAseq (scRNAseq) was performed using the 10X Genomics platform. Single-cell suspensions were counted using a hemocytometer and adjusted to 1,000 cells/μL. Sample processing through 10X Genomics and NextSeq 500 (Illumina) sequencing was performed by the Washington University Genome Technology Access Center (gtac.wustl.edu). Raw base call files were analyzed using Cell Ranger v.3.0.2. The filtered counts data from all 6 samples were combined in Partek Flow (www.partek.com). Samples were filtered on total reads, detected features, and mitochondrial content per cell to remove cell doublets and apoptotic debris. Unexpressed features were removed, and samples were normalized and log transformed. Principal component analysis was used to reduce dimensionality, and the first 20 principal components were further analyzed by graph-based (Louvain) clustering and t-SNE. The resulting 24 clusters were visualized in a two-dimensional t-SNE representation and were annotated to known biological cell types using canonical marker genes. The following cell types were annotated (selected markers are listed): Lymphocytes (CD3D, CD3G, IL7R); macrophages (C1QA, C1QB, C1QC); dendritic cells (CD1C and lack of expression of C1QA, C1QB and C1QC); retinal cells (SYNGR1, NPTX1); photoreceptors (GNAT1, GNGT1); RPE(PAX6, KRT8, KRT18); melanoma (DCT, TYRP1, PMEL).

### Mice

All experiments were performed using the NOD scid gamma (NOD.Cg-Prkdc^scid^ Il2rg^tm1Wjl^/SzJ: NSG) mice purchased from The Jackson Laboratory (#005557). All animal experiments were performed under protocols approved by the Animal Studies Committees of Washington University in St. Louis School of Medicine and Drexel University College of Medicine. FR was prepared from a 15 mM stock solution in DMSO at doses of 0.1, 0.3, 1.0, or 3.0 mg/kg and dextrose at a final volume of 200 μL. FR was administered subcutaneously every other day. Animals were randomly grouped (n=10/group) for placebo (1% DMSO) or FR treatments. MP41, MP46 and OCM1A cells were inoculated subcutaneously into the flanks of 5-week-old male NSG mice. Cells were prepared at a concentration of 2 x 10⁶ cells / 100 μL in a solution of ice-cold PBS / Matrigel Matrix (50/50 v/v) (Fisher Scientific # CB40234A). Primary tumors were allowed to grow to ~30 mm^2^ basal area based on measurements taken with calipers and using the formula (L × 2W)/2 where L and W are the longest and the shortest basal diameters of the tumor, respectively, before beginning treatment with FR. Tumors reaching a maximum length of 2 cm were defined as censored endpoints. At censored endpoints or at the end of each timecourse, mice were euthanized, and the tumors were removed using sterile surgical technique. Excised tumors were weighed to determine endpoint mass and snap-frozen in liquid nitrogen for later use.

### Immunoblotting

For tumor samples, approximately 100 mg of tumor was homogenized on ice in 1X Cell Lysis Buffer (Cell Signaling Technology, catalog# 9803) supplemented with 1 mM PMSF. For cultured cells, cells were grown in 10 cm dishes and treated with DMSO or 1 μM FR for 18 hr, and then lysed in 1X Cell Lysis Buffer. Lysates were sonicated on ice for 2 min (30 s on, 30 s off, 60% A), rotated end-over-end for 30 min, and cleared by centrifugation at 16,000 X g for 15 min. Protein concentration was determined using Bio-Rad Protein Assay Dye Reagent (Bio-Rad, catalog# 5000006). 15 μg of tumor protein was resolved on 12% SDS-PAGE gels and transferred to Immobilon-FL PVDF membrane (Millipore, catalog# IPFL00010). Membranes were blocked with 5% (w/v) BSA in TBST [20 mM Tris pH 7.6, 137 mM NaCl, 0.1% v/v Tween 20] and incubated with primary antibodies [Phospho-p44/42 MAPK (Erk1/2) Cell Signaling Technology, catalog# 4370S lot#24 and p44/42 MAPK (Erk1/2) Cell Signaling Technology catalog# 9107S lot#10]. Membranes were washed with TBST at least three times and incubated with IRDye 680–coupled goat anti-rabbit (LI-COR, catalog# 926-68071, lot# C90618-09) and IRDye 800 goat anti-mouse (LI-COR, catalog# 926-32210, lot# C91210-09 antibodies (LI-COR Biosciences). After incubation, membranes were washed at least three times with TBST, and signals were detected using Odyssey model 9120 imaging system (LI-COR Biosciences).

### Mouse radiotelemetry

Blood pressure and heart rate responses to chronic administration of vehicle or FR900359 were monitored by radiotelemetry in conscious mice by following previously described procedures (24, 51). Briefly, adult NSG male mice were implanted with a radiotelemetry pressure-sensing catheter in the right carotid artery under isoflurane (3% mixed with 95% oxygen) anesthesia. The body of the transmitter (HD-X10, Data Science International (DSI), St. Paul, MN) was tunneled into a subcutaneous pouch on the left flank of the animal. The mice were allowed to recover from the surgery for one week after which baseline blood pressure and heart rate were recorded for 24 hr, followed by alternating days of subcutaneous administration of vehicle or FR. After the 24-hour baseline recordings, the mice were randomly assigned to two groups: one group receiving FR (0.3mg/kg in dextrose, s.c.) and the other group receiving equal amounts of vehicle (2 μL DMSO in 200 μl of 5% dextrose, s.c.). Daily recordings were conducted from 1 – 5 pm for 14 days, and FR or vehicle was administered every other day at 3 pm. Continuous 24-hour BP recordings were conducted during the first injection and seventh injection of FR. Data was acquired and analyzed using DSI Ponemah software version 6.5. Averages of continuous systolic blood pressure (SBP), diastolic blood pressure (DBP), mean blood pressure (MBP), and heart rate (HR) recordings were plotted in one-hour intervals over 24 hours.

### Mouse behavioral tests

Mice were moved to the animal facility and allowed to acclimate to the new environment for 1 week prior to behavioral testing. All behavioral testing was conducted during the light cycle, by a female experimenter blinded to experimental group. All equipment was cleaned with 2% chlorhexidine diacetate or 70% ethanol between animals.

#### Open Field Activity/Exploratory Behavior

General activity levels and exploratory behavior was quantified over a 60-min period in an open-field (47.6 cm L × 25.4 cm W × 20.6 cm H) constructed of Plexiglas and surrounded by computerized photobeam instrumentation (Kinder Scientific, LLC, Poway, CA). General activity variables (total ambulations, rearings, time at rest) along with measures of emotionality, including time spent, distance traveled, and entries made into the central zone were analyzed.

#### Sensorimotor battery

Walking initiation, ledge, platform, pole, and inclined and inverted screen tests were performed as previously described (52). Time in each task was manually recorded. The average of two trials was used for analyses. Test duration was 60s, except for the pole test, which was extended to 120s. For walking initiation, time for an animal to leave a 21×21 cm^2^ square on a flat surface was recorded. For ledge and platform tests, the time the animal was able to balance on an acrylic ledge (0.75 cm wide and 30 cm high), and on a wooden platform (1.0 cm thick, 3.0 cm in diameter and elevated 47 cm) was recorded, respectively. The pole test was used to evaluate fine motor coordination by quantifying time to turn 180° and climb down a vertical pole. The screen tests assessed a combination of coordination and strength by quantifying time to climb up or hang onto a mesh wire grid measuring 16 squares per 10 cm, elevated 47 cm and inclined (60° or 90°) or inverted.

### Statistical Analyses

All statistical analyses were performed using IBM SPSS software (v.24) and Graphpad Prism 8. For cell line data, all experiments were performed in triplicate at least three times on different days. Means and standard errors were computed from all cell line data and t-tests were used to determine significance. For behavioral data, all data were screened for fit of distributions with assumptions underlying univariate analyses, which included the Shapiro-Wilk test and q-q plot investigations for normality. Means and standard errors were computed for each measure. Repeated measures Analysis of variance (ANOVA), and independent samples t-tests were used to analyze behavioral data. Statistical results were confirmed with two-tailed non-parametric testing, when available, for any datasets with violations of the univariate assumptions. Probability value for all analyses was *p* < .05, unless otherwise stated.

## Supporting information

Supplemental RNAseq data

## Acknowledgements

This work was supported by NIH grants GM124093 and CA234533 to K.J.B., GM118171 to J.A.C., and HL139754 to P.O. and by an AHA grant 16SDG2726027 to P. O. We thank Drs. Philip L. Custer and Steven M. Couch for surgical collection of enucleation samples. We thank the Alvin J. Siteman Cancer Center at Washington University School of Medicine and Barnes-Jewish Hospital in St. Louis, MO., for the use of the Siteman Flow Cytometry Core. The Siteman Cancer Center is supported in part by an NCI Cancer Center Support Grant #P30 CA091842. We thank the Genome Technology Access Center in the Department of Genetics at Washington University School of Medicine for help with genomic analyses. The Center is partially supported the Siteman Cancer Center and by ICTS/CTSA (UL1RR024992) from the National Center for Research Resources (NCRR), a component of the National Institutes of Health (NIH), and NIH Roadmap for Medical Research.

## Data and materials availability

Sequencing data will be made available on the GEO server for bulk RNAseq and scRNAseq experiments. All other data associated with this study are presented in the main text or supplementary materials.

