## Supplemental Data for "Targeting primary and metastatic uveal melanoma with a G protein inhibitor"

### Supplementary Data

#### Fig. S1

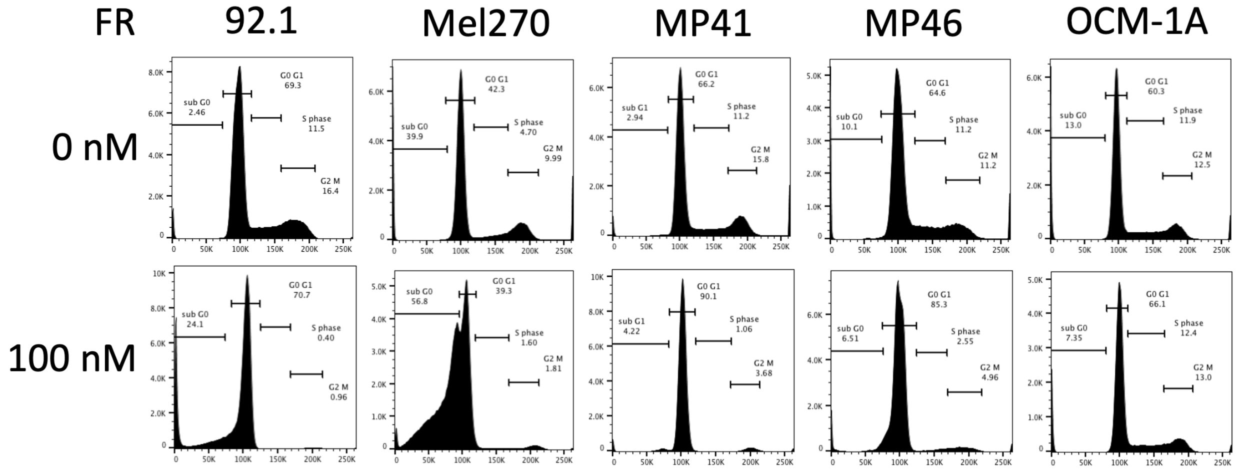


**Supplementary Figure S1.** DNA content histograms from cell cycle analysis of FR treated UM cell lines. Representative graphs showing number of cells (y-axis) versus intensity of propidium iodide staining for DNA content (x-axis) for 92.1 (GqQ209L), Mel270 (GqQ209P), MP41 (G11Q209L), MP46 (GqQ209L:BAP1null), and OCM-1A (BRAFV600E) UM cell lines. Top row – vehicle treated; bottom row – FR treated. Gating for cell cycle phases was done on vehicle treated histograms and applied to FR-treated histograms.

Fig. S2

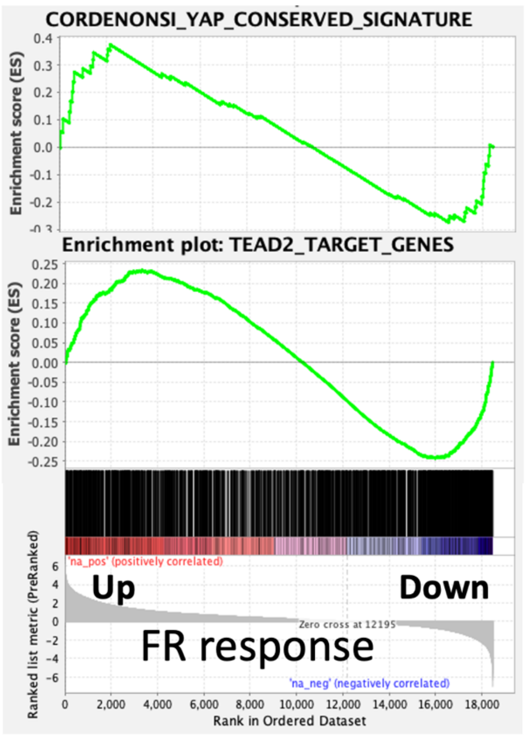


**Supplementary Figure S2.** Gene set enrichment analysis of YAP pathway in FR treated patient samples. Gene set enrichment analysis for YAP and TEAD target genes was performed using bulk RNAseq data from all FR-responsive human tumor samples. Upregulated and downregulated genes were enriched equally.

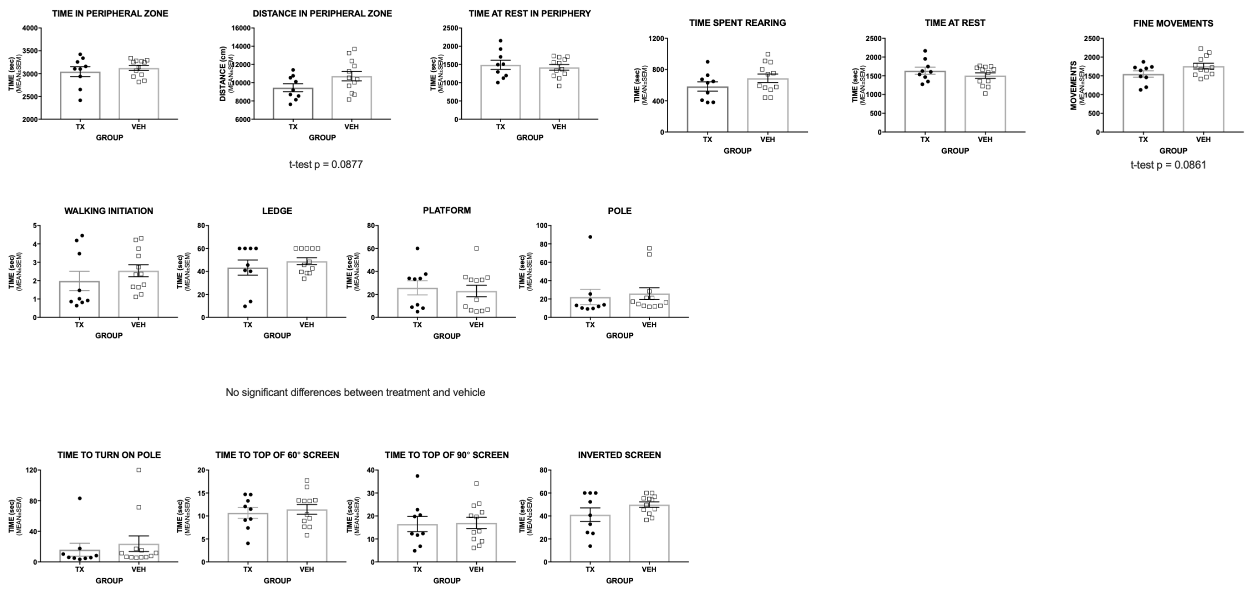


**Supplementary Figure S3.** Individual graphs of mouse behavior data in response to FR. Graphs correspond to summary data presented in Table 1. General activity levels and exploratory behavior were quantified over a 60-min period in an open-field, and general activity variables (total ambulations, rearings, time at rest) along with measures of emotionality, including time spent, distance traveled, and entries made into the central zone were analyzed. Time in walking initiation, ledge, platform, pole, and inclined and inverted screen tests were manually recorded. The average for two trials was used for each analysis. No significant differences ($p < 0.05$) were detected between control and FR-treated groups.

Fig. S4

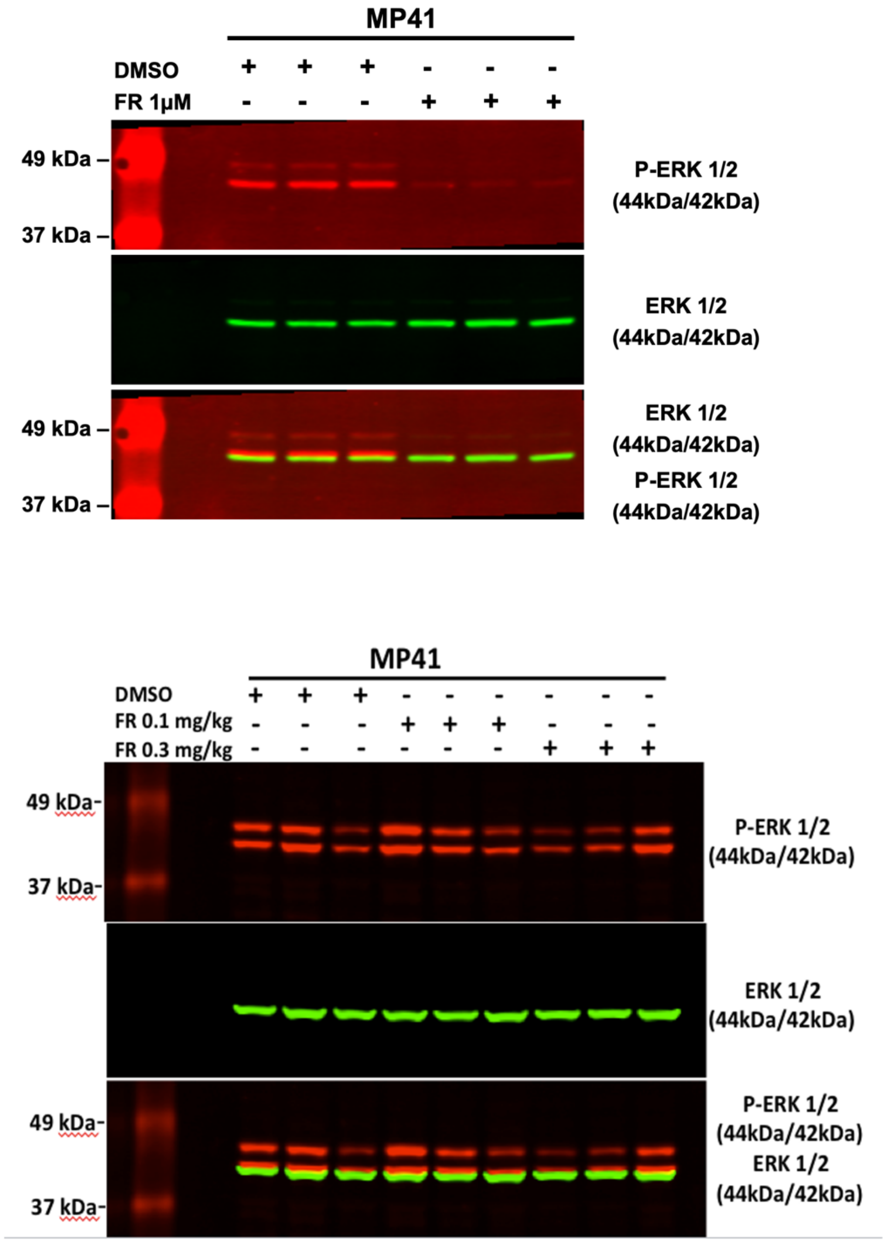


**Supplementary Figure S4.** Immunoblots for phospho-ERK and total ERK on cells and tumor lysates. Cultured cells were treated with vehicle (DMSO) or 1 μM FR for 18 hr; approximately 100 mg of xenograft tumor was collected from treated mice. Immunoblots were probed with primary antibodies against phospho-p44/42 MAPK (P-ERK1/2) and p44/42 MAPK (ERK1/2) followed by IRDye–coupled secondary antibodies. Signals were detected using Odyssey model

9120 imaging system (LI-COR Biosciences). Panels show individual and merged fluorescent signals for P-ERK1/2 (top) and total ERK1/2 (middle) from a single representative immunoblot.

**Table S5:** Human tumor samples used for RNAseq analysis

| | Molecular Class | GNAQ | GNA11 |
|---|---|---|---|
| UM030 | 1 | Q209P | wt |
| UM031 | 2 | Q209L | wt |
| UM032 | 1 | wt | wt |
| UM033 | 1 | Q209L | wt |
| MUM04 | nd | Q209L | wt |
| UM043 | 1 | wt | Q209L |
| UM044 | 2 | wt | Q209L |
| UM045 | 1 | wt | Q209L |
| UM046 | 2 | wt | Q209L |
| UM047 | 1 | Q209P | wt |
| UM049E | 2 | wt | Q209L |
| UM051E | 2 | wt | Q209L |
| UM052E | 2 | wt | Q209P |

**Supplementary Table S5.** Human samples used for RNAseq analyses. Ten fine-needle aspiration biopsy samples and three enucleation samples were collected from UM patients as described in Methods. Molecular classes were determined by Castle Biosciences. GNAQ and GNA11 mutations were determined from the RNAseq data. Molecular classification was not performed for the liver metastasis sample MUM04.

**Supplementary Data S6. (.xlsx)** List of genes showing significant response to FR by RNAseq. ENSEMBL identifiers, Genbank names and brief descriptions are given for each gene. Data in columns are log2 fold change and p-value (two-tailed, paired T-test) based on bulk RNAseq cpm data for all nine FR-responsive samples or broken out into Class 1 and Class 2 sample subsets.
